# Zinc Fingers and Homeoboxes 2 is Required for Diethylnitrosamine-induced Liver Tumor Formation in C57BL/6 Mice

**DOI:** 10.1101/2022.09.02.506400

**Authors:** Jieyun Jiang, Courtney Turpin, Guofang (Shirley) Qiu, Mei Xu, Eun Lee, Terry D. Hinds, Martha L. Peterson, Brett T. Spear

## Abstract

Liver cancer, comprised mainly of hepatocellular carcinoma (HCC), is the third leading cause of cancer deaths worldwide and increasing in Western countries. We previously identified the transcription factor Zinc Fingers and Homeoboxes 2 (Zhx2) as a regulator of hepatic gene expression, and many Zhx2 target genes are dysregulated in HCC. Here, we investigate HCC in Zhx2-deficient mice using the Diethylnitrosamine (DEN)-induced liver tumor model. Our study using whole-body Zhx2 knock-out (*Zhx2*^*KO*^) mice revealed the complete absence of liver tumors 9 and 10 months after DEN exposure.Analysis soon after DEN treatment showed no differences in expression of the DEN bioactivating enzyme CYP2E1 and DNA polymerase delta 2, or in the numbers of γH2AX foci between *Zhx2*^*KO*^ and wild-type (*Zhx2*^*wt*^) mice. The absence of Zhx2, therefore, did not alter DEN bioactivation or DNA damage. *Zhx2*^*KO*^ livers showed fewer positive foci for Ki67 staining and reduced IL-6 and AKT2 expression compared to *Zhx2*^*wt*^ livers, suggesting that Zhx2 loss reduces liver cell proliferation and may account for reduced tumor formation. Tumors were reduced but not absent in DEN-treated liver-specific Zhx2 knock-out mice, suggesting that Zhx2 acts in both hepatocytes and non-parenchymal cells to inhibit tumor formation. Analysis of data from The Cancer Genome Atlas and Clinical Proteomic Tumor Consortium indicated that ZHX2 mRNA and protein levels were significantly higher in HCC patients and associated with clinical pathological parameters.

**Conclusions:** In contrast to previous studies in human hepatoma cell lines and other HCC mouse models showing that Zhx2 acts as a tumor suppressor, our data indicate that Zhx2 acts as an oncogene in the DEN-induced HCC model and is consistent with the higher ZHX2 expression in HCC patients.

## INTRODUCTION

Liver cancer is the sixth most common cancer and the third leading cause of cancer deaths worldwide (1). Hepatocellular carcinoma (HCC), which accounts for 85%-90% of primary liver cancer, is increasing dramatically in Western countries due, in large part, to the continued rise of obesity-associated non-alcoholic fatty liver disease (NAFLD) (2). NAFLD, with a globally estimated frequency of 25% (3), can progress to non-alcoholic steatohepatitis (NASH), which includes the development of inflammation and fibrosis (4), which, if left untreated, can worsen to cirrhosis and HCC (4, 5). With the alarming rise in obesity in the United States (US), the incidence of both NAFLD and HCC is also expected to grow in coming years (5, 6); by 2030, liver cancer is projected to be the third leading cause of cancer deaths in the US (7). HCC is also among the most lethal cancers, with a five-year survival rate of 5-8%, which is attributed to both late-stage diagnosis and few treatment options (8). Liver transplantation is one option, but the need far exceeds the number of donors (9). Sorafenib and Regorafenib, the only approved drugs for HCC, have limited efficacy and extend survival by an average of fewer than three months (10), and numerous other pharmaceutical compounds have failed to meet clinical end points in Phase III trials (11). Thus, an improved understanding of HCC development and additional HCC biomarkers are needed to monitor and combat this increasingly prevalent cancer.

A hallmark of many cancers, including HCC, is the reactivation of genes that are normally expressed during fetal developed and silenced at birth. Alpha-fetoprotein (AFP), which is abundantly expressed in the fetal liver, was the first HCC-associated “oncofetal” gene identified (12). This has led to considerable interest in AFP regulation, including a study showing that postnatal AFP serum levels were higher in BALB/cJ mice than any other strain (13).The persistent AFP expression in BALB/cJ was transmitted as a monogenic trait (13). By positional cloning, we showed that this trait was due to a hypomorphic mutation in the BALB/cJ *Zinc fingers and homeoboxes 2* (*Zhx2*) gene that dramatically reduced Zhx2 levels (14, 15). Zhx2 is a member of a small family that also contains Zhx1 and Zhx3, all of which contain two C2-H2 zinc fingers motifs and four or five homeodomains (16). Several studies indicate that Zhx2 is a transcriptional regulator, although the mechanisms by which Zhx2 controls target genes are not known and a consensus Zhx2 binding site has not been identified (16).

In addition to AFP, several additional HCC-associated oncofetal genes, including Glypican 3 (*Gpc3*), lipoprotein lipase (*Lpl*) and the long non-coding RNA H19, continue to be expressed in the adult BALB/cJ liver. More recently, we developed C57BL/6 mice in which a floxed Zhx2 gene has been deleted in hepatocytes. AFP, Gpc3, Lpl, and H19 continue to be expressed in Zhx2-deficient adult livers similarly to BALB/cJ mice, confirming that this trait is due to the absence of Zhx2 (14, 17, 18)

Dysregulation of oncofetal genes in the absence of Zhx2 in the postnatal liver has led us to propose that Zhx2 could function as a tumor suppressor to control liver tumorigenesis. Human clinical data in support of this notion are conflicting. Several studies suggest that Zhx2 levels are increased in HCC whereas others indicate reduced Zhx2 levels (19, 20). Tissue culture and xenograft studies using human hepatoma cell lines indicated that Zhx2 inhibits cell proliferation and reduces levels of Cyclins A and E (21), supporting the possibility that Zhx2 functions as a tumor suppressor. Liver-specific Zhx2 knock-out mice had increased tumors compared to wild-type controls after hydrodynamic tail-vein injection (HTVI) of the oncogenes AKT and Myc (22), consistent with tumor suppressor activity of Zhx2.

To further investigate the mechanisms by which Zhx2 impacts HCC in mice, we used the well-established Diethylnitrosamine (DEN) model of HCC (23). Here, 14 day-old wild-type (*Zhx2*^*wt*^) and whole-body Zhx2 knock-out (*Zhx2*^*KO*^) mice were given a single injection of DEN. Upon bioactivation, toxic DEN metabolites can damage cell components, including DNA, resulting in the appearance of HCC tumors after 8-9 months. Both male and female *Zhx2*^*wt*^ mice treated with DEN exhibited numerous tumors, with the tumor burden being greater in males than in females. Although we expected greater tumor incidence in *Zhx2*^*KO*^ mice based on the predicted Zhx2 function as a tumor suppressor, our data surprisingly showed that DEN-treated *Zhx2*^*KO*^ mice had a complete absence of tumors. Analysis of livers within a week after DEN treatment indicates that this absence could be due to decreased cell proliferation in *Zhx2*^*KO*^ livers. Furthermore, tumors were reduced, but not absent, when the DEN treatment was performed in liver-specific Zhx2 knock-out mice, suggesting that Zhx2 acts in both hepatocytes and non-parenchymal cells (NPCs) to inhibit tumor formation after DEN treatment.UALCAN analysis of data from The Cancer Genome Atlas (TCGA) and Clinical Proteomic Tumor Consortium (CPTAC) indicated that ZHX2 mRNA and protein levels were significantly higher in HCC patients, suggesting that our results may provide insight into the role of ZHX2 in HCC progression in humans.

## EXPERIMENTAL PROCEDURES

### Experimental Animals

All mice used in this study were on the C57BL/6 background and were housed in the University of Kentucky Division of Laboratory Animal Research (DLAR) facility in accordance with Institutional Animal Care and Use Committee-approved protocols. To generate Zhx2 knock-out mice, exon 3, which encodes the entire Zhx2 coding region, was flanked by loxP sites to generate a Zhx2 floxed allele (*Zhx2*^*fl*^) (24). The *Zhx2*^*fl*^ mice were crossed with Alb-Cre mice (Jackson, cat.# 003574) or E2a-Cre mice (Jackson, cat.# 003724) to generate liver-specific Zhx2 knock-out mice (*Zhx2*^*Δliv*^) or whole-body Zhx2 knock-out mice (*Zhx2*^*KO*^), respectively (25).

### Tumor Induction and Analysis

For the DEN-induced HCC model, *Zhx2*^*KO*^ and *Zhx2*^*wt*^ littermates were given a single intraperitoneal injection of DEN (Sigma-Aldrich) [(10 mg/kg), diluted in Phosphate-buffered saline (PBS)] or vehicle (PBS) at 14 days of age, weaned and maintained on a regular chow diet. Tumor loads were examined at 9 or 10 months. Externally visible tumors (≥0.5 mm) were counted and measured. Partial large lobes were fixed in formalin, and paraffin-embedded sections were processed for Hematoxylin and Eosin (H&E) staining. Remaining lobes were microdissected into tumors and non-tumor tissues and stored at −80°C. For short-term studies, *Zhx2*^*KO*^ and *Zhx2*^*wt*^ littermates were given DEN (10 mg/kg) or vehicle (PBS) at 14 days of age and killed at designated timepoints within a week after DEN exposure. H&E stained liver sections were used for histopathological evaluation of HCC by Eun Lee in a blinded manner.

### Immunohistochemistry (IHC) and Quantitative Image Analysis

IHC staining was performed on paraffin-embedded tissue sections. Detailed methods and antibodies used for IHC can be found in Supplemental Materials.

Aperio ScanScope XT high-throughput digital slides scanner system (Aperio, Vista, CA, USA) was used to image entire stained slides at 20x magnification to create a single high-resolution digital image.

Quantification of nuclear staining for Ki67 was done using HALO imaging software (Indica Labs, Albuquerque, NM, USA).

### Real-Time Quantitative PCR (RT-qPCR)

Total RNAs were extracted from samples with RNAzol RT reagent (Molecular Research Center) according to the manufacturer’s instructions. One microgram of RNA was reverse-transcribed to cDNA using the High-Capacity cDNA Reverse Transcription Kit (ThermoFisher Scientific). RT-qPCR reactions were prepared with SsoAdvanced Universal SYBR Green Supermix (Bio-Rad) and amplified in a Bio-Rad CFX96 real-time PCR system. Oligonucleotides (Supplemental Table 1) were obtained from Integrated DNA Technologies (Coralville, IA). The qPCR Ct values were normalized to ribosomal protein L30 or GAPDH levels and reported as the normalized expression of the indicated gene using the ΔCt method (26).

### Western Blot analysis

For western blotting, liver or tumor protein lysates were prepared and quantified. Each sample was resolved by electrophoresis using 10% SDS-PAGE and transferred to PVDF membranes or Immobilon-FL membranes. After incubation with corresponding primary antibodies overnight at 4°C and the corresponding secondary antibodies, protein expression was visualized. Detailed methods and antibodies used for western blotting can be found in Supplemental Materials.

### Statistical Analysis

Statistical analyses were performed using GraphPad Prism 9.0.2 software. All values within a group were averaged and plotted as mean ± SEM. One-way or two-way ANOVA or Student’s *t* test were used to assess statistical significance. A *p* value less than or equal to 0.05 was considered significant.

### UALCAN Analysis

The UALCAN database (http://ualcan.path.uab.edu) was used to analyze RNA-seq data from The Cancer Genome Atlas (TCGA) (27) and protein expression data from the Clinical Proteomic Tumor Consortium (CPTAC) (28). ZHX2 protein expression values from the CPTAC data portal were log2 normalized in each sample. Then a Z-value for each sample was calculated as standard deviations from the median across samples (28). ZHX2 mRNA expression in TCGA Liver Hepatocellular Carcinoma (LIHC) samples was compared to normal samples. LIHC samples were then categorized using clinical patient data from TCGA-LIHC project as follows: 1) Tumor histology; 2) Individual cancer stages: based on American Joint Committee on Cancer pathologic tumor stage information, samples were divided into stage I, stage II, stage III and stage IV group; 3) Tumor grade: where tumor grade information is available, samples were categorized into grade 1, grade 2, grade 3, and grade 4 groups (27).

## RESULTS

### Zhx2 expression is effectively eliminated in the livers of Zhx2 knock-out mice

We previously described C57BL/6J mice with a floxed *Zhx2* allele (Figure 1A, *Zhx2*^*fl*^; (24)); Zhx2 levels are the same in *Zhx2*^*fl*^ mice as in mice with the wild-type Zhx2 gene (*Zhx2*^*wt*^; data not shown). We also showed that breeding *Zhx2*^*fl*^ mice to albumin-Cre to delete Zhx2 in hepatocytes (*Zhx2*^*Δliv*^) or βactin-Cre to delete Zhx2 in all cells (*Zhx2*^*KO*^) dramatically reduced or completely eliminated liver Zhx2 mRNA levels, respectively (25). To confirm changes in Zhx2 protein levels in knock-out mice, immunohistochemical staining was performed using adult liver from *Zhx2*^*wt*^, *Zhx2*^*KO*^, and *Zhx2*^*Δliv*^ mice. Zhx2 was present in the nuclei of both hepatocytes and non-parenchymal cells in *Zhx2*^*wt*^ mice (Figure 1B). Hepatocytes from *Zhx2*^*Δliv*^ mice no longer expressed Zhx2, although protein was still detected in non-parenchymal cells, whereas Zhx2 was not detected in any cells of *Zhx2*^*KO*^ liver (Figure 1B). This data confirms the complete absence of Zhx2 in *Zhx2*^*Δliv*^ hepatocytes and Zhx2 expression in non-parenchymal cells.

**Figure 1.**
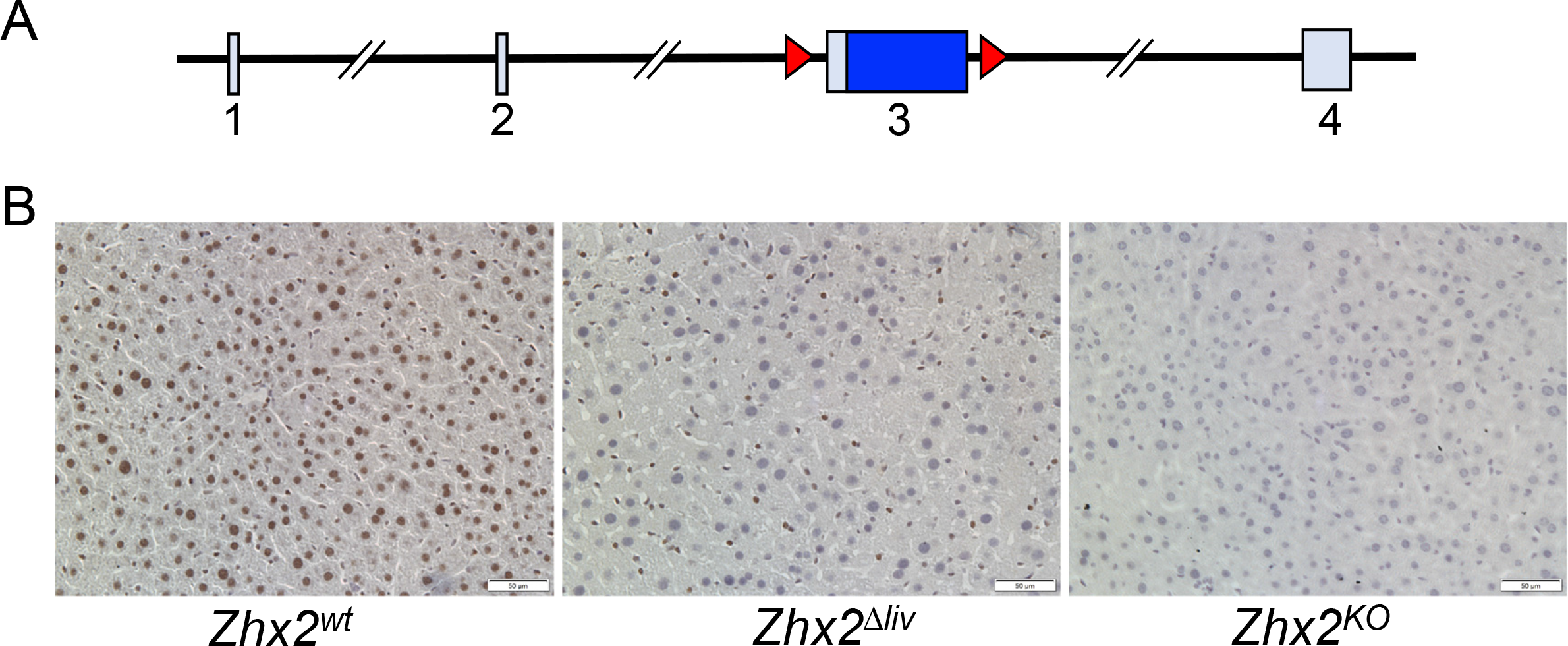
The *Zhx2* gene is expressed in hepatocytes and non-parenchymal cells and efficiently deleted in knock-out mouse livers. ***A***. The mouse *Zhx2* gene spans 145 kb and is comprised of 4 exons, including a 2.7 kb exon 3 that encodes the entire 836 amino acid of Zhx2 protein; *loxP* sites (red triangles) used for deletion of flanked exon 3. ***B***. *IHC staining of adult livers with anti-Zhx2*. Adult mouse liver sections from homozygous *Zhx2*^*wt*^ mice, *Zhx2*^*Δliv*^ mice (*Zhx2*^*fl/fl*^, *Alb-Cre*^*+*^), and *Zhx2*^*KO*^ mice (*Zhx2*^*fl/fl*^, *βactin-Cre*^*+*^) were stained with Zhx2 antibodies and counterstained with Hematoxylin. [Magnification 20×]

### Zhx2 is essential for HCC formation after DEN treatment in male and female mice

Previous studies using hepatoma cell lines indicated that increased Zhx2 expression resulted in reduced cell proliferation *in vitro* and reduced xenograft growth in nude mice (21) and that HTVI of oncogenes AKT/Myc led to more tumors in Zhx2 liver-knock-out mice (22), consistent with Zhx2 functioning as a tumor suppressor. To explore the role of Zhx2 in the initiation and progression of tumor development *in vivo*, we used the well-established DEN model in *Zhx2*^*wt*^ and *Zhx2*^*KO*^ mice. The whole-body *Zhx2*^*KO*^ mice were used since any Zhx2-expressing cells could potentially contribute to a liver tumor phenotype. Mice were injected with DEN or PBS on postnatal day 14 (p14) and killed at 9 months after DEN exposure, at which time livers were removed and analyzed. No tumors were observed in cohorts that received PBS injections (Table 1, Figure 2A, B). Tumors were present in 100% (14/14) of the *Zhx2*^*wt*^ male mice and 25% (3/12) of *Zhx2*^*wt*^ female mice. The tumor incidence and tumor number were less in female mice, consistent with previous studies (29). In stark contrast to *Zhx2*^*wt*^ mice, visible liver tumors were completely absent in all male (0/11) and female (0/10) *Zhx2*^*KO*^ mice. To test whether there was a delay in tumor formation, DEN-treated *Zhx2*^*wt*^ and *Zhx2*^*KO*^ male and female mice were killed at 10 months after p14 DEN treatment (for ethical reasons, DEN-treated *Zhx2*^*wt*^ male mice could not be maintained longer than this time). While maximal tumor size and tumor numbers increased in *Zhx2*^*wt*^ male mice (Figure 2B) and tumor incidence increased to 36% (4/11) in *Zhx2*^*wt*^ female mice (Table 1), no visible tumors were present in *Zhx2*^*KO*^ male and female livers. These data demonstrate that Zhx2 is required for HCC progression after perinatal DEN treatment and indicate that Zhx2 promotes tumor growth in this model.

**Table 1.** Tumor incidence in *Zhx2*^*KO*^ mice and *Zhx2*^*wt*^ littermates after DEN exposure.

|  | Treatment | Tumor Incidence |  |  |  |
| --- | --- | --- | --- | --- | --- |
|  |  | Male |  | Female |  |
|  |  | *Zhx2<sup>wt</sup>* | *Zhx2<sup>KO</sup>* | *Zhx2<sup>wt</sup>* | *Zhx2<sup>KO</sup>* |
| 9 months | PBS | 0% (0/15) | 0% (0/9) | 0% (0/14) | 0% (0/10) |
|  | DEN | 100% (14/14) | 0% (0/11) | 25% (3/12) | 0% (0/10) |
| 10 months | PBS | 0% (0/10) | 0% (0/11) | 0% (0/9) | 0% (0/10) |
|  | DEN | 100% (10/10) | 0% (0/11) | 36% (4/11) | 0% (0/11) |

*Zhx2<sup>KO</sup>* mice and *Zhx2<sup>wt</sup>* littermates were given a single intraperitoneal injection of DEN (10 mg/kg, Sigma-Aldrich) or vehicle (PBS) at 14 days of age. Livers were removed and tumor loads were examined 9 or 10 months after DEN exposure. Number of mice with tumor/cohort in parentheses.

**Figure 2.**
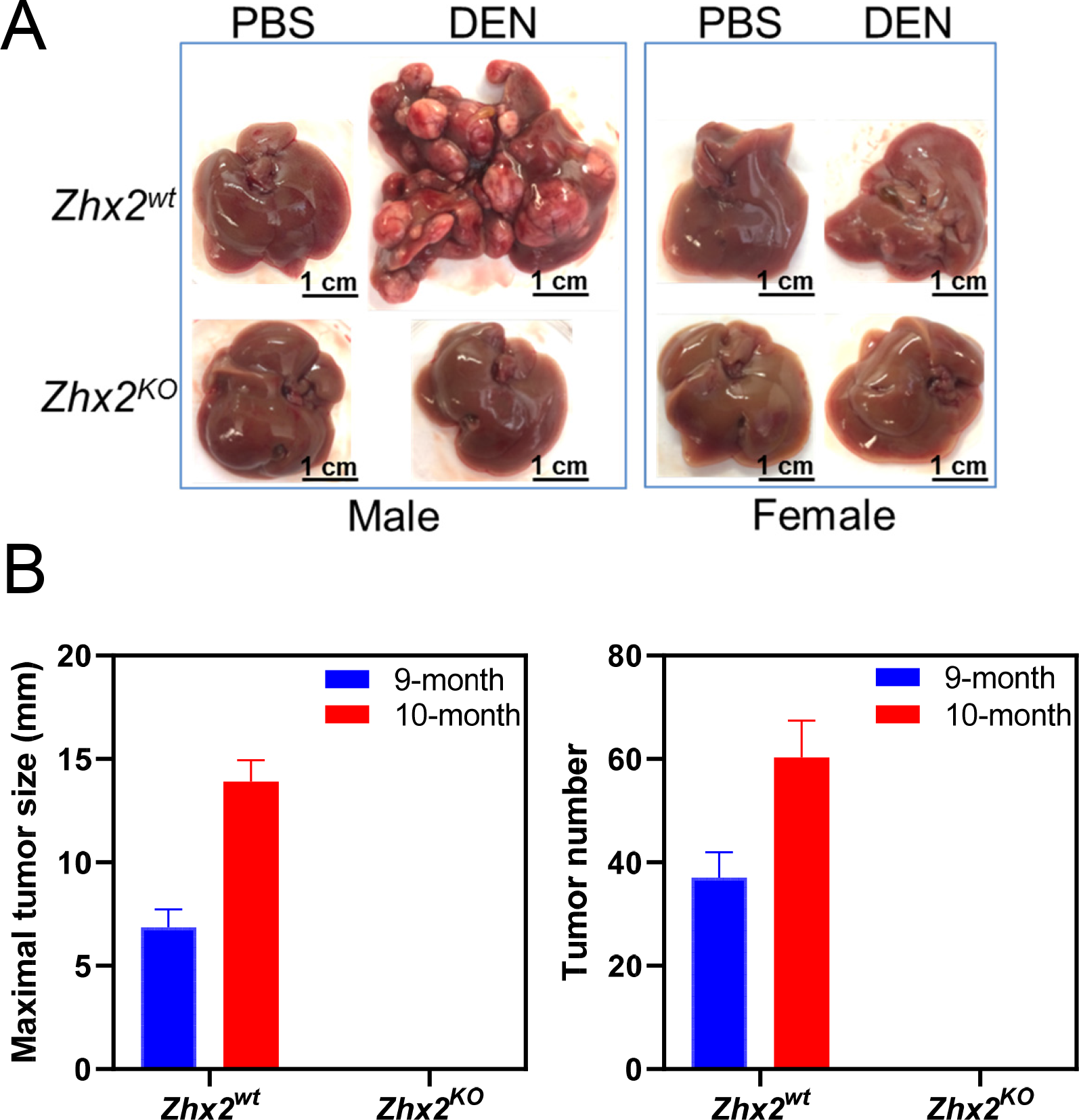
The absence of Zhx2 in *Zhx2*^*KO*^ mice completely blocks DEN-induced HCC. ***A***. Male and female *Zhx2*^*wt*^ and *Zhx2*^*KO*^ mice (9∼15 mice/cohort) were given a single i.p. injection DEN (10 mg/kg, Sigma-Aldrich) or vehicle (PBS) at p14. After nine months, livers were removed, and tumors were quantitated. Representative livers are shown; tumor incidence data are shown in Table 1. ***B***. The maximal tumor size (left panel) and the average number of visible tumors (right panel) in *Zhx2*^*wt*^ male mice at nine months (n= 14) and ten months (n=10) and *Zhx2*^*KO*^ male mice at nine months (n=11) and ten months (n=11) after DEN treatment.

H&E stained liver sections from DEN-treated *Zhx2*^*wt*^ male mice confirmed that tumors were HCC (Figure 3A) and also confirmed the absence of tumors or any small foci in livers of DEN-treated *Zhx2*^*KO*^ male mice or any of the PBS-treated control mice. Previous studies indicated that DEN-initiated tumors are Glutamine Synthetase (GS)-negative, which was confirmed in tumors in *Zhx2*^*wt*^ mice; the absence of Zhx2 did not alter GS staining after PBS or DEN treatment (Figure 3B).

**Figure 3.**
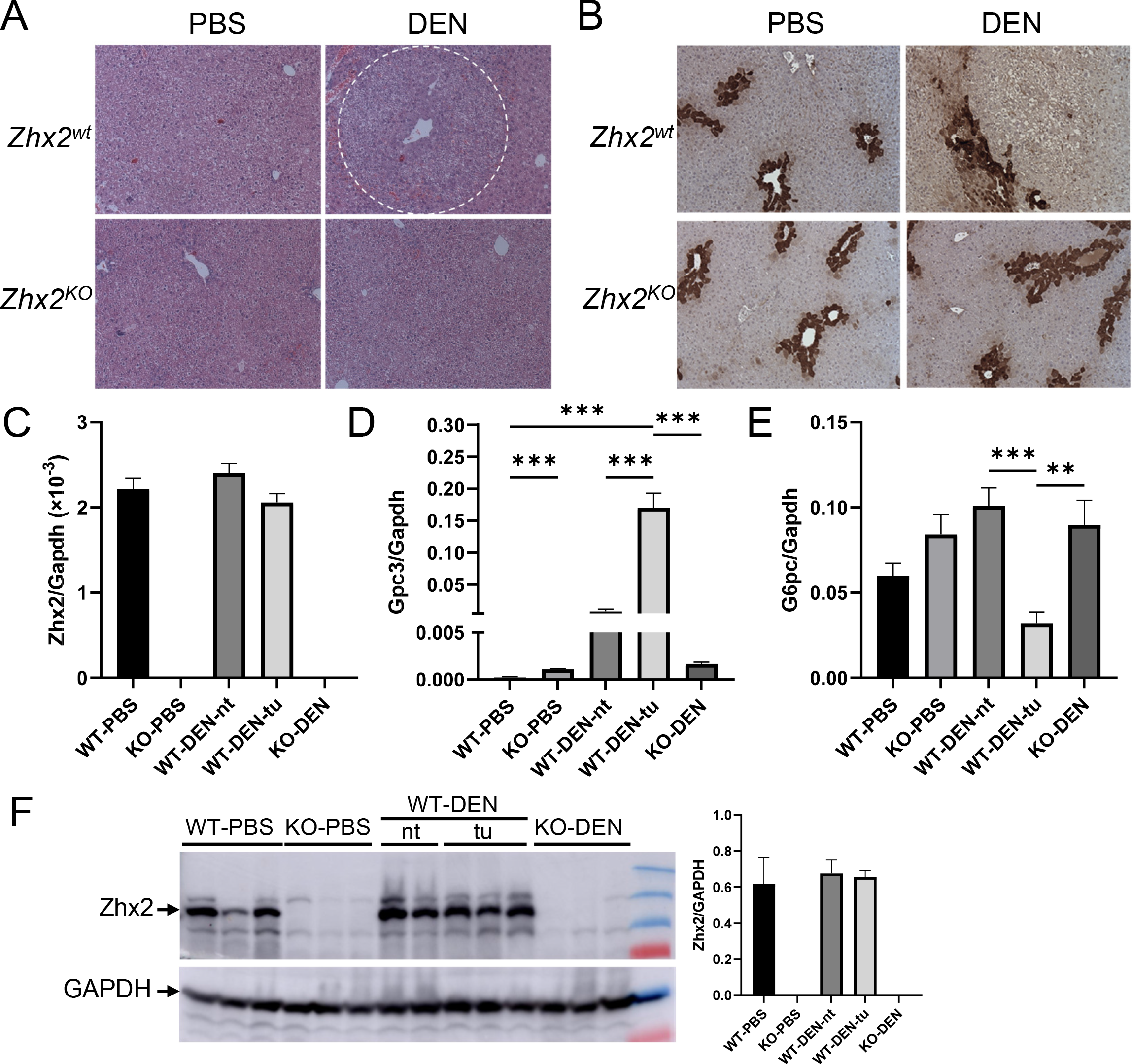
Histological and mRNA analysis of livers nine months after PBS or DEN treatment confirm the presence of HCC *Zhx2*^*wt*^ mice and the absence of tumors in *Zhx2*^*KO*^ mice. ***A, B***. H&E staining (A) and immunohistochemical staining for Glutamine synthetase (B) in *Zhx2*^*wt*^ and *Zhx2*^*KO*^ male mice after PBS or DEN treatment. ***C, D, E***. Quantitation of hepatic Zhx2 (C), Gpc3 (D) and G6PC (E) mRNA levels by RT-qPCR in PBS- or DEN-treated *Zhx2*^*KO*^ mice (n=8 and 10, respectively) or livers from PBS-treated *Zhx2*^*wt*^ mice (n=9) or tumors (tu) or non-tumor tissues (nt) from DEN-treated *Zhx2*^*wt*^ mice (n=10). mRNA levels were normalized to GAPDH. **, *p*<0.01; ***, *p*<0.001. ***F***. Zhx2 protein levels were analyzed by Western blot of liver lysate from PBS- or DEN-treated *Zhx2*^*KO*^ mice or livers from PBS-treated *Zhx2*^*wt*^ mice or tumors (tu) or non-tumor tissues (nt) from DEN-treated *Zhx2*^*wt*^ mice. Blots were re-probed with antibodies against GAPDH. The Zhx2 protein levels were quantified using ImageQuant™ TL Image Analysis Software (GE Healthcare) and normalized to GAPDH.

Gpc3, one of many oncofetal genes controlled by Zhx2, has emerged as an important HCC biomarker in humans (18, 30). We therefore analyzed Zhx2 and Gpc3 levels in the 9-month liver samples. As expected, Zhx2 mRNA and protein were not detected in PBS- or DEN-treated *Zhx2*^*KO*^ livers (Figure 3C, F). In contrast, Zhx2 was expressed at similar levels in PBS-treated *Zhx2*^*wt*^ livers and in both non-tumor regions and tumors present in DEN-treated *Zhx2*^*wt*^ livers, indicating that Zhx2 mRNA and protein levels are not different in HCC (Figure 3C, F). Gpc3 mRNA levels were barely detectable in PBS-treated *Zhx2*^*wt*^ livers and roughly 5-fold higher in PBS-treated *Zhx2*^*KO*^ livers, consistent with previous studies showing the incomplete silencing of Gpc3 after birth in the absence of Zhx2. Hepatic Gpc3 levels were dramatically higher in both non-tumor and tumor tissue of DEN-treated *Zhx2*^*wt*^ mice compared to PBS-treated *Zhx2*^*wt*^ livers; however, this increase was significantly greater in HCC tumors. In contrast, Gpc3 mRNA levels in DEN-treated *Zhx2*^*KO*^ livers were similar to those of the PBS-treated *Zhx2*^*KO*^ controls; this similarity is consistent with the absence of tumor in DEN-treated *Zhx2*^*KO*^ livers. It has been reported that glucose-6-phosphatase (G6PC) is downregulated in DEN-induced pre-malignant lesions (31).When compared to PBS-treated controls, the G6PC mRNA level was significantly decreased in tumors of DEN-treated *Zhx2*^*wt*^ mice as seen previously but remained unchanged in *Zhx2*^*KO*^ livers (Figure 3E).

### Hepatic CYP2E1 levels and DNA damage are the same in *Zhx2*^*wt*^ and *Zhx2*^*KO*^ mice soon after DEN treatment

The striking absence of liver tumors in DEN-treated *Zhx2*^*KO*^ mice led us to focus on events associated with the initiation of tumorigenesis soon after DEN administration. DEN must be hydroxylated to a bioactive form that is toxic to cells and can damage DNA, both of which can lead to cellular transformation that will ultimately manifest in tumors (23). Since CYP2E1 is the primary enzyme for DEN bioactivation, we analyzed CYP2E1 levels in *Zhx2*^*wt*^ and *Zhx2*^*KO*^ livers prior to and up to 24 hours after DEN treatment. No difference in hepatic CYP2E1 mRNA levels was observed in mice with or without Zhx2 at 0, 4, 8, or 24 hours after DEN treatment (Figure 4A), although levels in both groups of mice increased slightly at 24 hrs. Western blot analysis indicated that CYP2E1 protein levels also were the same in *Zhx2*^*wt*^ and *Zhx2*^*KO*^ livers at all timepoints tested (Figure 4C and Supplemental Figure 1A). We also determined whether Zhx2 levels changed in response to DEN treatment. Western analysis confirmed the absence of Zhx2 in *Zhx2*^*KO*^ livers but indicated that Zhx2 protein levels did not change 24 hours after DEN treatment in *Zhx2*^*wt*^ mice (Figure 4D and Supplemental Figure 1B).

**Figure 4.**
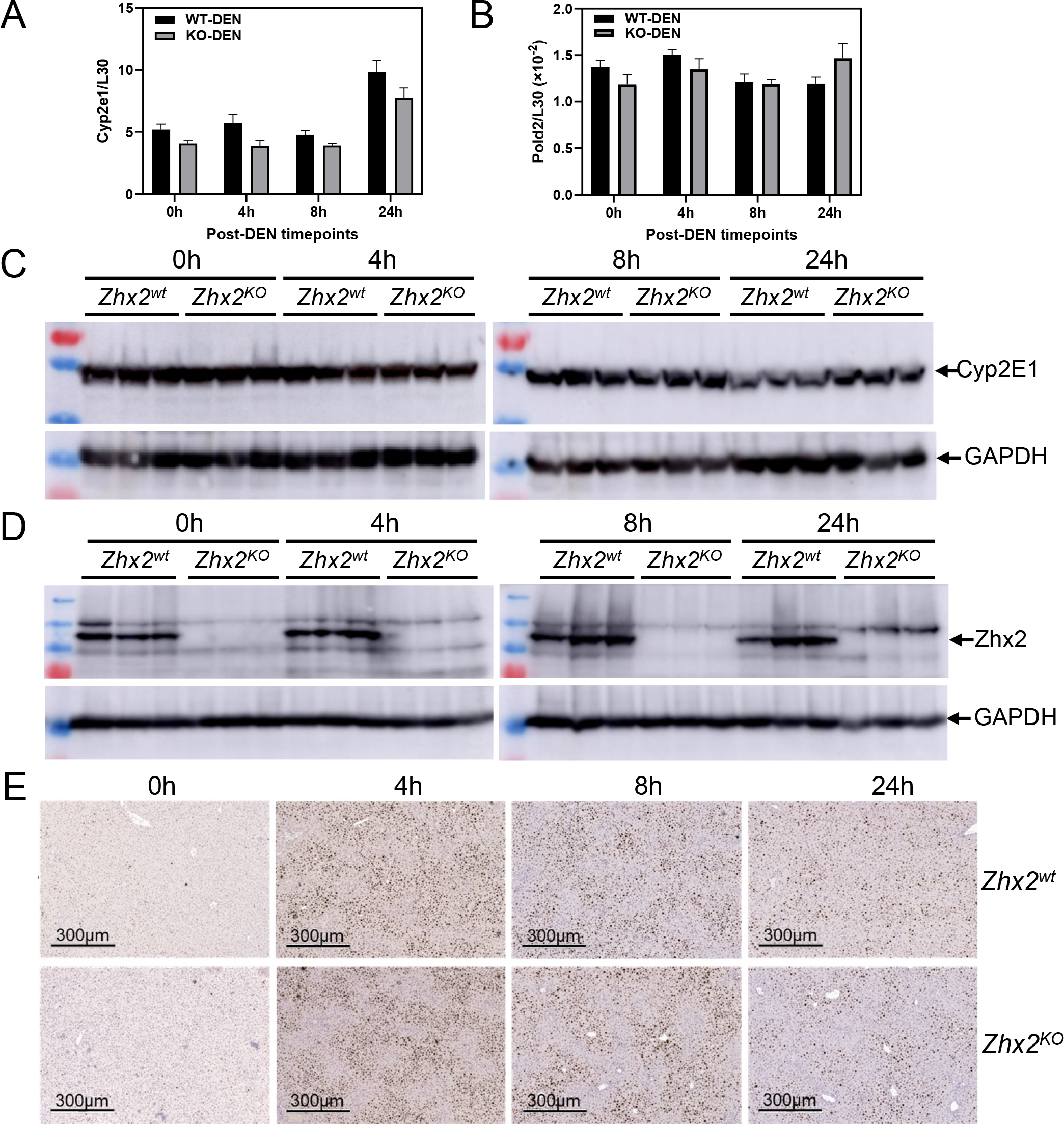
CYP2E1, Pold2 expression and γH2AX staining are the same in *Zhx2*^*wt*^ and *Zhx2*^*KO*^ mice 24 hours after DEN treatment. Data were obtained from p14 *Zhx2*^*wt*^ and *Zhx2*^*KO*^ mice prior to (0h) and 4, 8, or 24 hours after DEN treatment. ***A, B***. Hepatic Cyp2e1 (A) and Pold2 (B) mRNA levels were quantitated by RT-qPCR and normalized to ribosomal protein L30. ***C, D***. Hepatic CYP2E1 (C) and Zhx2 (D) protein levels were analyzed by Western blot; each lane represents an individual mouse. Blots were re-probed with antibodies against GAPDH. ***E***. Representative images from liver sections after immunohistochemical staining for γH2AX and counterstained with Hematoxylin.

Bioactive DEN damages DNA causing a rapid increase in double-strand breaks that can be detected by γH2AX staining (32). Therefore, we analyzed γH2AX staining in *Zhx2*^*wt*^ and *Zhx2*^*KO*^ livers. Prior to DEN treatment, very little staining was observed in either *Zhx2*^*wt*^ or *Zhx2*^*KO*^ livers. Staining increased dramatically four hours after treatment, indicating a large number of double-strand breaks, with a gradual reduction by 24 hours. However, there was no difference in γH2AX staining between the two cohorts (Figure 4E). Expression of the four members of the DNA polymerase delta (Pold) family, which are involved in DNA repair after damage, were also measured after DEN treatment. No difference in mRNA levels of Pold2 (Figure 4B) or other Pold family members (data not shown) was observed between *Zhx2*^*wt*^ and *Zhx2*^*KO*^ between 0 and 24 hours after DEN treatment. Taken together, this data indicates that differences in DNA damage or DNA repair soon after DEN treatment are unlikely to explain the lack of tumors in *Zhx2*^*KO*^ livers.

### Hepatic cell proliferation is decreased in *Zhx2*^*KO*^ mice compared to *Zhx2*^*wt*^ soon after DEN treatment

Although DEN metabolites can act as genotoxins, DEN is also hepatotoxic and triggers an inflammatory response that results in activation of NPCs and elevated expression of interleukin-6 (IL-6), a mitogen which promotes compensatory proliferation of surviving hepatocytes; this proliferation plays a critical role in DEN-induced hepatocaracinogenesis (29, 33). This led us to test whether proliferation was affected by the absence of Zhx2 after DEN exposure. Livers from p14 *Zhx2*^*wt*^ and *Zhx2*^*KO*^ mice were removed prior to DEN treatment or at several timepoints up to one week after treatment, and sections were stained for Ki67. Similar Ki67 staining was seen in untreated livers from both cohorts (Figure 5A, B). The number of Ki67-positive nuclei was slightly lower in DEN-treated *Zhx2*^*KO*^ livers than in *Zhx2*^*wt*^ livers at 8 hours and 1 day after DEN treatment. However, at 2 days after DEN treatment, there are dramatically fewer Ki67-positive nuclei in *Zhx2*^*KO*^ livers compared to the *Zhx2*^*wt*^ livers (Figure 5A, B).After seven days, numerous Ki67-positive nuclei were still present in *Zhx2*^*wt*^ liver sections whereas very few positive cells were seen in *Zhx2*^*KO*^ liver sections. RT-qPCR data showed hepatic IL-6 mRNA levels were significantly decreased in *Zhx2*^*KO*^ livers compared to *Zhx2*^*wt*^ livers at 2- and 7-days post-DEN treatment, whereas hepatic TNFα mRNA levels remain unchanged between *Zhx2*^*KO*^ livers and *Zhx2*^*wt*^ livers at all time points 7 days post-DEN treatment (Figure 5C). These results suggest that decreased cell proliferation may be a consequence of reduced IL-6 expression in NPCs of *Zhx2*^*KO*^ livers. This, in turn, could account for the lack of tumors after DEN treatment in *Zhx2*^*KO*^ mice. The reduced proliferation rates are analogous to the lower phosphorylation of AKT (pAKT) observed in the *Zhx2*^*KO*^ mice compared to the *Zhx2*^*wt*^ (Figure 5D). We further analyzed the AKT expression in DEN-treated livers. We found that AKT2 expression was significantly lower in the DEN-treated *Zhx2*^*KO*^ mice compared to *Zhx2*^*wt*^ littermates but there was no change in AKT1 levels (Figure 5E, F). NF-κB, a downstream effector of AKT, was lower, although this difference did not reach statistical significance (Figure 5E, F). These results suggest that reduced AKT2 expression and lower pAKT are associated with reduced proliferation rates in the *Zhx2*^*KO*^ mice compared to *Zhx2*^*wt*^ littermates after DEN treatment.

**Figure 5.**
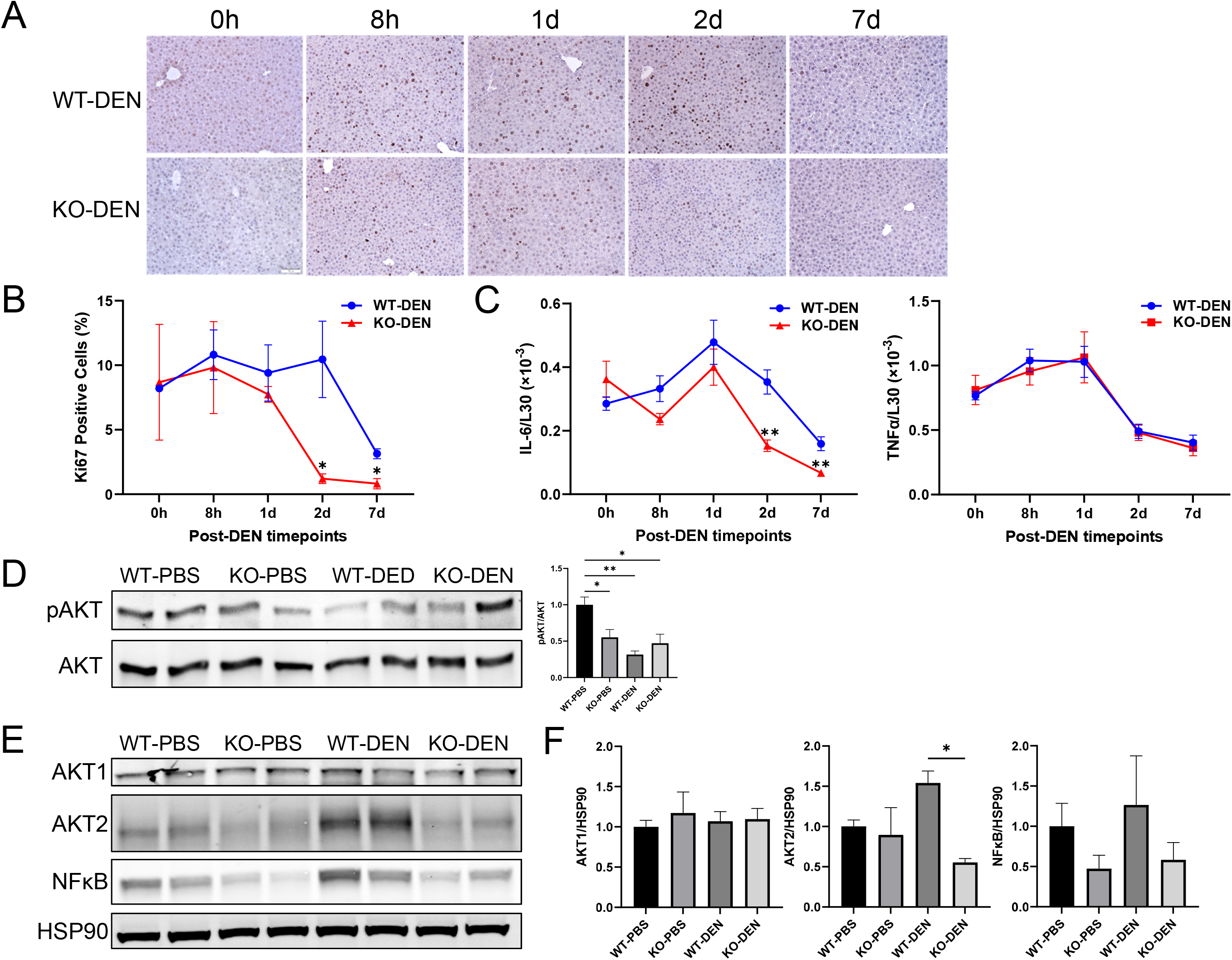
Cell proliferation is reduced in the livers of *Zhx2*^*KO*^ mice compared to *Zhx2*^*wt*^ mice during the seven days after DEN treatment. ***A***. Liver sections were obtained from p14 mice prior to (0h) and 8h, 1d, 2d, and 7d after DEN treatment and stained for the proliferation marker Ki67. Representative images from liver sections at designated timepoints. ***B***. Quantitation of Ki67 IHC staining. Aperio ScanScope XT digital slides scanner (Aperio, Vista, CA, USA) was used to scan the entire stained slide at 20x magnification to create a single high-resolution digital image. Quantification of nuclear staining for Ki67 was done using HALO imaging software (Indica Labs, Albuquerque, NM, USA). Data were from three mice in each group. *, *p*<0.05. ***C***. Hepatic IL-6 and TNFα mRNA levels were quantitated by RT-qPCR and normalized to ribosomal protein L30. **, *p*<0.01. ***D-F***. Immunoblotting and densitometry of p-AKT (D), AKT1, AKT2, NF-κB p65 and heat shock protein 90 (HSP90) (E, F) from the 7d livers of PBS- or DEN-treated *Zhx2*^*KO*^ mice or *Zhx2*^*wt*^ mice. *, *p*<0.05, **, *p*<0.01 (n=4).

### Zhx2 expression in both hepatocytes and non-parenchymal cells contributes to reduced tumor formation after DEN treatment

As noted previously, Zhx2 is expressed in both hepatocytes and non-parenchymal liver cells. Although DEN is metabolized in hepatocytes, it is possible that the absence of Zhx2 in other liver cell types could explain the lack of tumors after DEN treatment. To address this, DEN was injected into male p14 *Zhx2*^*Δliv*^ mice and control littermates. As shown previously (Figure 1), the loss of Zhx2 in the *Zhx2*^*Δliv*^ mice is restricted to hepatocytes. After nine months, mice were killed, and livers were analyzed. In contrast to the complete absence of tumors in *Zhx2*^*KO*^ livers, a small number of tumors were present in *Zhx2*^*Δliv*^ livers (Figure 6), although tumor numbers were considerably less than in *Zhx2*^*fl*^ livers.Furthermore, the maximal tumor size and the liver/body weight ratios were lower in *Zhx2*^*Δliv*^ mice than in *Zhx2*^*fl*^ mice. This data indicates that the absence of Zhx2 in both hepatocytes and non-parenchymal cells contributes to the reduced tumors after DEN treatment.

**Figure 6.**
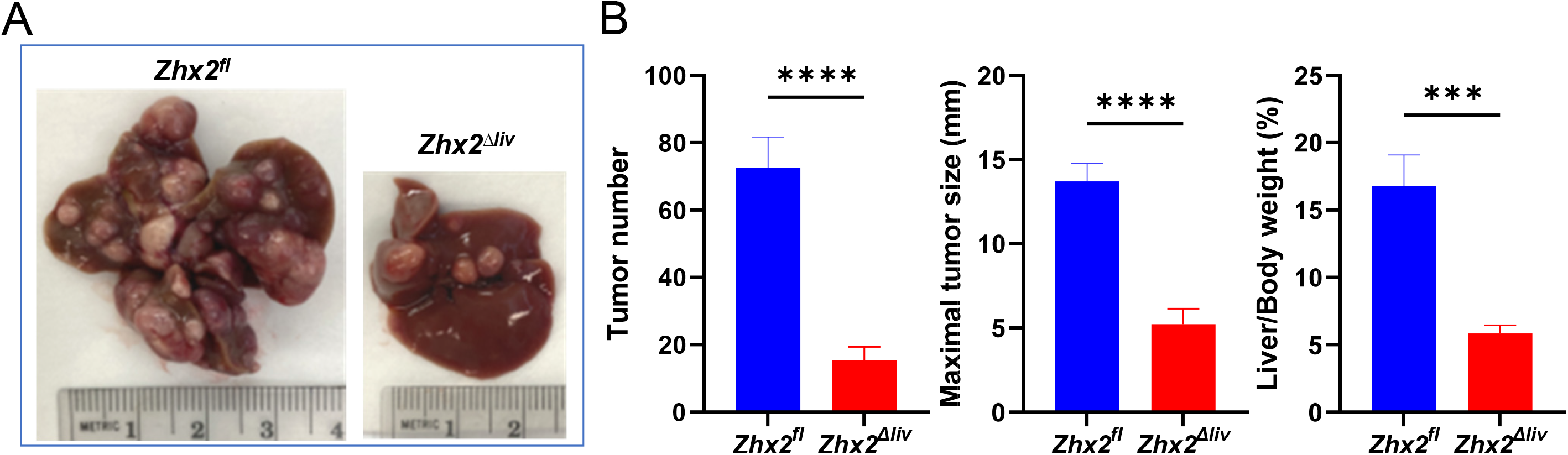
DEN-induced Tumors are significantly reduced in *Zhx2*^*Δliv*^ mice compared to *Zhx2*^*fl*^ littermate controls. ***A***. Representative livers from *Zhx2*^*fl*^ and *Zhx2*^*Δhep*^ mice nine months after DEN treatment. ***B***. Liver tumor number (left panel), maximal tumor size (middle panel), and liver/body weight ratio (right panel) of *Zhx2*^*fl*^ (n=7) and *Zhx2*^*Δliv*^ (n=9) mice nine months after DEN treatment. ***, *p*<0.001; ****, *p*<0.0001.

### ZHX2 mRNA and protein levels are significantly higher in HCC patients and associated with clinical pathological parameters

UALCAN can be used to compare relative gene expression and protein levels between normal and tumor samples, as well as between tumor subgroups stratified by pathological grade, clinical stage, age, sex, and other clinical features (27, 28). We used UALCAN to analyze the ZHX2 mRNA levels in the TCGA database and protein levels in the CPTAC database. ZHX2 protein levels were significantly higher in HCC samples compared to normal samples (Figure 7A). From TCGA-LIHC RNA-seq data, ZHX2 mRNA levels were significantly higher in LIHC samples than normal samples (Figure 7B). Based on histology subtype, ZHX2 mRNA levels were significantly higher in HCC (361 cases, the majority of 371 LIHC cases), fibrolamellar carcinoma (FLC, 3 cases) and hepatocholangiocarcinoma (HCC-CC, 7 cases) (Figure 7C). Also, ZHX2 mRNA expression significantly increased with the individual cancer stage 1 to 3 and the tumor grade 1 to 4 (Figure 7D, E). Expression in cancer stage 4 didn’t differ, possibly due to the low number of cases. Overall, these results showed that ZHX2 expression was significantly correlated with HCC tumor stage and grade.

**Figure 7.**
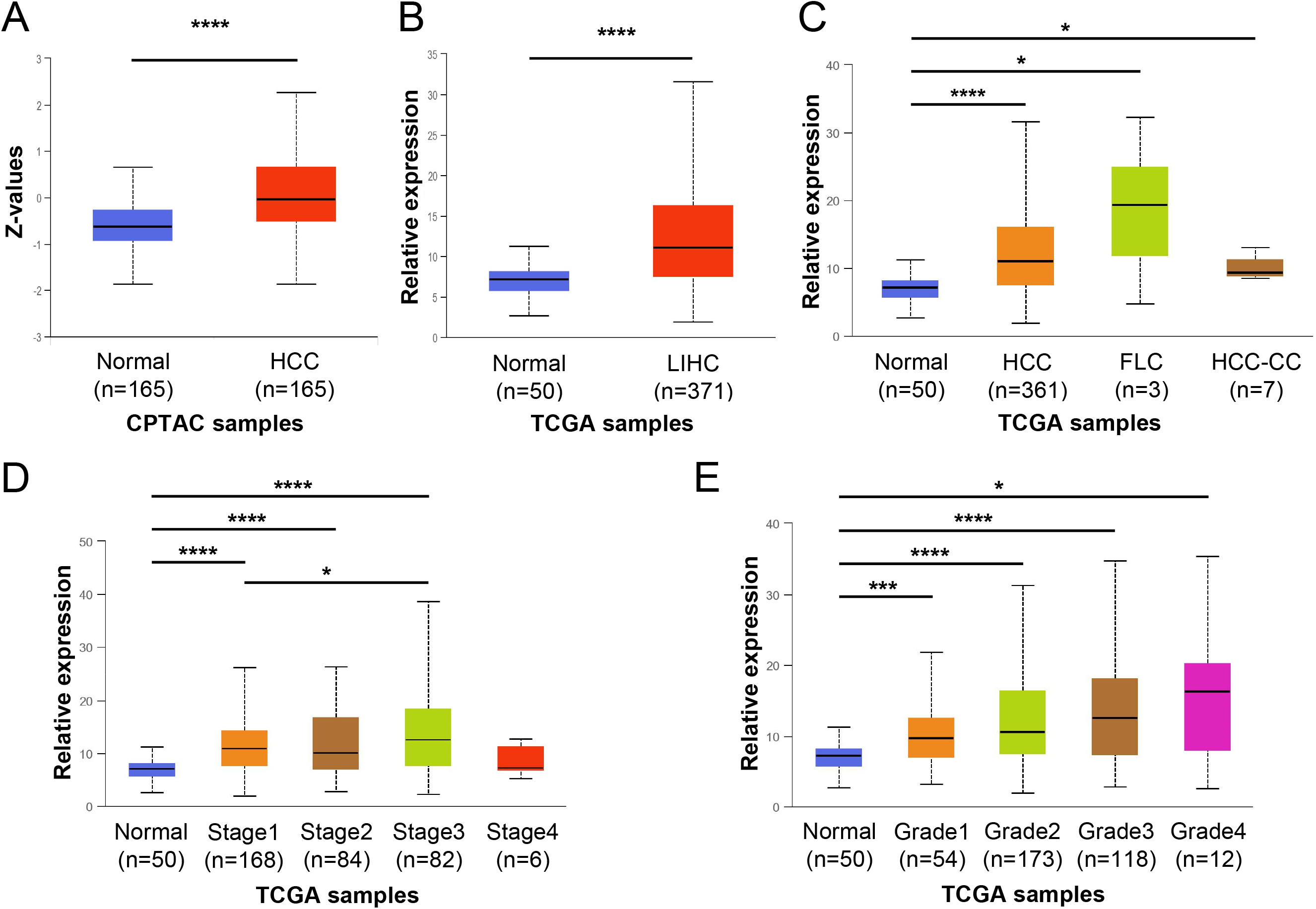
ZHX2 mRNA and protein levels are significantly higher in HCC patients and associated with clinical pathological parameters. UALCAN was used to analyze data from The Cancer Genome Atlas (TCGA) and the Clinical Proteomic Tumor Consortium (CPTAC). ***A***. ZHX2 protein expression in CPTAC samples. HCC, hepatocellular carcinoma. ***B***. ZHX2 mRNA expression in TCGA Liver Hepatocellular Carcinoma (LIHC) samples. ***C-E***. ZHX2 mRNA expression in TCGA-LIHC samples based on histological subtypes (C), individual cancer stages (D), and tumor grade (E). FLC, fibrolamellar carcinoma; HCC-CC, hepatocholangiocarcinoma. *, *p*<0.05; ***, *p*<0.001; ****, *p*<0.0001.

**Figure 8.**
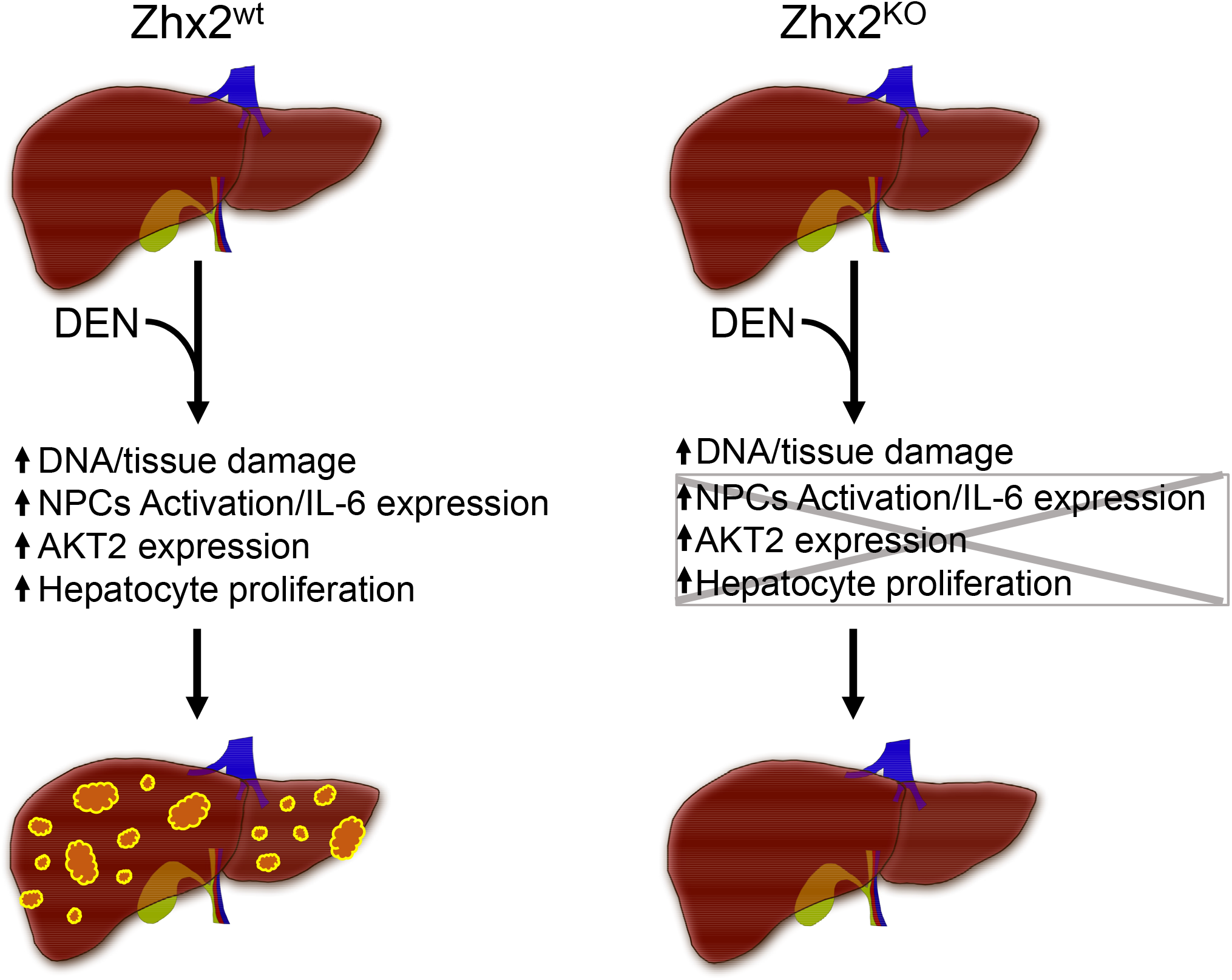
Graphic summary of action of Zhx2 in DEN-induced HCC. The absence of Zhx2 completely blocks liver tumor formation 9 months after DEN treatment. While DEN-mediated DNA damage is the same in both *Zhx2*^*KO*^ and *Zhx2*^*wt*^ mice, hepatocyte proliferation, AKT2 activation and IL-6 expression are reduced in *Zhx2*^*KO*^ livers soon after DEN exposure. These data suggest that Zhx2 functions in both hepatocytes and non-parenchymal cells (NPCs), a possibility that is supported by the fact that tumors are reduced, but not absent, in DEN-treated *Zhx2*^*Δliv*^ mice.

## DISCUSSION

Data presented here show that DEN-initiated hepatocarcinogenesis is completely blocked in the absence of Zhx2, with a complete lack of tumors at 9 months and 10 months of age in both male and female mice. H&E-stained sections and molecular analyses both support the absence of tumors in these mice. Most tellingly, Gpc3 levels dramatically increased in tumors (∼700-fold) of DEN-treated *Zhx2*^*wt*^ livers compared to PBS-treated *Zhx2*^*wt*^ livers. “Non-tumor” tissues from DEN-treated *Zhx2*^*wt*^ livers also had higher level of Gpc3 (33-fold), likely due to the presence of multiple smaller tumor nodules present in these tissues. However, Gpc3 levels were not increased in *Zhx2*^*KO*^ livers after DEN treatment compared to PBS-treated *Zhx2*^*KO*^ livers. In addition, when compared to PBS-treated controls, G6PC mRNA levels significantly decreased in tumors of DEN-treated *Zhx2*^*wt*^ mice as seen previously (31) but remained unchanged in DEN-treated *Zhx2*^*KO*^ livers.

Since differences between *Zhx2*^*wt*^ and *Zhx2*^*KO*^ mice soon after DEN treatment could explain the disparity in tumor formation, we focused on the first week after treatment. CYP2E1, the primary bioactivator of DEN in the liver to generate reactive radicals which can damage cellular components, including DNA, and lead to an inflammatory response, is present at similar levels in *Zhx2*^*wt*^ and *Zhx2*^*KO*^ livers. γH2AX staining and Pold mRNA levels were also similar between *Zhx2*^*wt*^ and *Zhx2*^*KO*^ livers during the early timepoints, suggesting that there was no difference in cellular or DNA damage and repair in these cohorts after DEN treatment. However, cell proliferation, as measured by Ki67 staining, was higher and persisted longer in *Zhx2*^*wt*^ mice than in *Zhx2*^*KO*^ mice during the week after DEN treatment. This increased proliferation could increase the likelihood of DNA damage and/or other tumor initiation events in *Zhx2*^*wt*^ mice that resulted in tumors 9 months later. The reduced proliferation rates align with the lower pAKT and AKT2 levels observed in the *Zhx2*^*KO*^ mice compared to the *Zhx2*^*wt*^. AKT2 expression has been correlated with HCC prognosis in humans (34), suggesting that targeting Zhx2 may have the potential to regulate this pathway for HCC therapy. How Zhx2 may regulate the AKT-pathway, by gene regulation or protein-protein interaction, needs further investigation. Targeting the AKT-pathway in HCC has been proposed (35). Possible concerns of antagonizing this pathway alone for HCC therapy may cause other deleterious effects such as inducing insulin-resistant diabetes.However, these effects have not been observed in the *Zhx2*^*KO*^ mice and implies that this could be a better and safer target. Knockout studies of the AKT isozymes in mice have shown positive results, as Akt1 ablation impaired carcinogenesis (36), and Akt2 deletion reduced the occurrence of HCC in c-Met-transfected mice (37). Future studies that dissects the AKT isozyme function in HCC and how each might impact growth or metabolic pathways is needed.

Our data that Zhx2 acts as a tumor promoter after DEN treatment is consistent with higher ZHX2 expression in human HCC from TCGA and CPTAC data and suggests that this model may provide insight into Zhx2 and HCC progression in humans. However, our mouse data appears paradoxical with other liver tumor models indicating that Zhx2 functions as a tumor suppressor. This suggests that the action of Zhx2 in liver cancer is context-dependent, a notion that is consistent with a growing number of studies of other tumor promoters and/or suppressors (38). For example, the tyrosine phosphatase Shp2 (38), Stat3 (38), beta-catenin (38), the microRNA miR-21 (39), and proteins in the NF-kB pathway (29) can all promote or inhibit tumor formation in different tumor models. Interestingly, the Zhx2 target H19 has also been shown to have both tumor-promoting and tumor-suppressing activities in different liver tumor models (40). In vitro growth and xenograft tumor studies suggested that Zhx2 represses the proliferation of liver tumor cell lines, which may be due to Zhx2-mediated repression of cyclin A and cyclin E expression (21). In contrast, we found that cell proliferation, as measured by Ki67 staining, was greater in the presence of Zhx2. Thus, differences in the control of cell proliferation by Zhx2 in transformed liver cell lines and perinatal hepatocytes could at least partially explain the seemingly contradictory results in these distinct model systems and help explain the conflicting data that has been obtained in human studies (19, 20).

A recent study by Zhao et al. showed that Zhx2 alleviates NASH through PTEN activation (41). This study utilized a high-fat high-cholesterol diet to induce NASH and used both liver-specific Zhx2 knock-out and liver-specific Zhx2 transgenic mice. Mice in this study did not develop tumors, so we cannot compare this model with our DEN model regarding HCC formation. However, it should be noted that Zhao et al. (41) and Yue et al (21) focused on hepatocytes, although it is increasing clear that NPCs also play critical roles in NAFLD progression from simple steatosis to NASH, cirrhosis and HCC (42). DEN-induced HCC depends on an inflammatory response and IL-6 production in activated Kupffer cells (29, 33). Our data showed that hepatic IL-6 levels were significantly decreased in whole-body Zhx2 knock-out mice, which might due to the absence of Zhx2 in NPCs. Thus, understanding Zhx2 function in both NPCs and hepatocytes will be necessary to fully understand the role of Zhx2 in HCC initiation and progression, and could help elucidate the basis for the contradictory data with Zhx2 in different liver disease models and may provide valuable insight into the complexity of HCC progression.

Many Zhx2 target genes are dysregulated in HCC, and it is possible that one or more of these target genes is responsible for the dramatic difference in HCC progression between *Zhx2*^*wt*^ and *Zhx2*^*KO*^ mice after DEN treatment. Several known Zhx2 target genes, including H19, Gpc3 and Lpl, have been associated with liver cancer. H19, which acts as a tumor suppressor specifically in the DEN model (43), could help explain the lack of tumors in *Zhx2*^*KO*^ livers since it is expressed at higher levels in the absence of Zhx2. A majority of reports indicate that Gpc3 is pro-oncogenic (30) and Lpl levels were higher in human and mouse HCC (44). However, hepatic Gpc3 and Lpl levels are higher in *Zhx2*^*KO*^ than in *Zhx2*^*wt*^ mice, which argues that these proteins are not involved in the absence of tumors in DEN-treated *Zhx2*^*KO*^ mice. Altered lipid metabolism is found in HCC, including DEN-induced tumors (45).Many enzymes regulating lipid homeostasis are controlled by Zhx2 (46, 47), so changes in hepatic lipids could also contribute to tumor differences between *Zhx2*^*wt*^ and *Zhx2*^*KO*^ livers.

Zhx2 is expressed in hepatocytes and non-parenchymal cells in the adult liver (Figure 2, (25)). The fact that tumors are reduced in DEN-treated *Zhx2*^*Δliv*^ mice, compared to the absence of tumors in DEN-treated *Zhx2*^*KO*^ mice, indicates that non-parenchymal cells contribute to tumor formation. BALB/cJ mice, which have a natural Zhx2 mutation, have reduced atherosclerotic lesions compared to mice with a wild-type Zhx2 gene when placed on a high fat diet. These BALB/cJ mice have greater numbers of anti-inflammatory M2 macrophages, which could account for reduced damage compared to Zhx2-positive mice with higher pro-inflammatory M1 macrophages (46). Based on these data, Zhx2 expression in Kupffer cells and/or infiltrating macrophages in *Zhx2*^*Δliv*^ mice compared to *Zhx2*^*KO*^ mice could lead to different inflammatory environments after DEN treatment that would impact tumorigenesis. Future studies will need to investigate the role of Zhx2 in various liver cell populations in relation to liver disease, including HCC.

While early Zhx2 studies focused on Zhx2 function in HCC due to its control of AFP and other hepatic genes, a growing number of reports have associated Zhx2 with other cancers. Zhx2 levels are low in Hodgkin’s Lymphoma and Multiple Myeloma, suggesting that Zhx2 may function as a tumor suppressor in these B cell malignancies although functional studies were not performed (48). Zhx2 inhibits the metastatic potential of several thyroid cancer cell lines, possibly by inhibition of S100A14, suggesting a tumor suppressor role in this cancer (49). In contrast to these tumors, Zhx2 is oncogenic in clear cell Renal Cell Carcinoma, the most common kidney tumor, potentially through co-regulation of target genes with NF-κB (50), and in triple-negative breast cancer, where it co-regulates target genes with HIF1a (51). Future studies will require different liver tumor model systems to fully understand Zhx2 control of tumor initiation and promotion. Since Zhx2 is known to regulate many genes that control lipid homeostasis, and that obesity and NAFLD are currently major drivers of the increased incidence of HCC, it will be particularly interesting to investigate the role of Zhx2 in high-fat diet-induced liver tumor models.

## List of abbreviations

HCC: hepatocellular carcinoma
Zhx2: Zinc Fingers and Homeoboxes 2
DEN: diethylnitrosamine
NAFLD: non-alcoholic fatty liver disease
NASH: non-alcoholic steatohepatitis
AFP: Alpha-fetoprotein
Gpc3: Glypican 3
Lpl: Lipoprotein lipase
G6PC: Glucose-6-phosphatase
GS: Glutamine Synthetase
CYP2E1: Cytochrome P450 enzyme 2E1
Pold2: DNA polymerase delta 2
PBS: Phosphate-buffered saline
*Zhx2*^*fl*^: Zhx2 floxed allele
*Zhx2*^*wt*^: Zhx2 wild-type
*Zhx2*^*KO*^: Zhx2 knock-out
*Zhx2*^*Δliv*^: Zhx2 liver-specific knock-out
TCGA: The Cancer Genome Atlas
CPTAC: Clinical Proteomic Tumor Consortium
IL-6: interleukin-6.

## SYNOPSIS

We found the absence of Zhx2 completely blocks liver tumor formation in mice after DEN treatment. Our data indicates that reduced cell proliferation and AKT2 expression could result in lower tumors. The reduced IL-6 expression in *Zhx2*^*KO*^ mice suggests that non-parenchymal cells also contribute to the lack of tumors, possibly by reducing compensatory hepatocyte proliferation after DEN-mediated damage.

These data indicate that Zhx2 is pro-oncogenic in the DEN-induced mouse HCC model and is consistent with the higher expression of ZHX2 in HCC patients.

## AUTHORS CONTRIBUTIONS

J.J. and B.T.S. conceived and designed the experiments. J.J., C.T., S.Q. performed the experiments. M.X. performed immunoblotting and calculations for Figure 5D-5F. E.L. evaluated histopathology of liver H&E sections. J.J. and B.T.S. analyzed data, wrote and edited the manuscript. M.L.P. and T.H. contributed to discussions of experimental design and interpretation and manuscript editing. All authors have read and approved the final version of the manuscript.

## ACKNOWLEDGMENTS

Floxed Zhx2 mice were obtained from the trans-NIH Knock-Out Mouse Project.

## DATA AVAILABILITY

The authors confirm that the data supporting the findings of this study are available within the article.

## Supplemental materials include

**Supplemental Experimental Procedures**

**Supplemental Table 1**

**Supplemental Table 2**

**Supplemental Figure 1**

## Supplemental Experimental Procedures

### Immunohistochemistry (IHC)

Immunohistochemical (IHC) staining was performed on paraffin-embedded tissue sections. Staining of Zhx2 was performed using rabbit anti-Zhx2 (Abcam, ab205532) and Dako EnVision System HRP Kit (Catalog# K4011) following the manufacturer’s protocol. Other IHC staining used antibodies against Phospho-Histone H2AX (γH2AX) (Cell Signaling, #9718), Ki67 (Fisher Scientific, Catalog no. 50-245-563), or Glutamine Synthetase (Santa Cruz Biotechnology, sc-74430). Briefly, antigen retrieval using High pH IHC Antigen Retrieval Solution (Invitrogen, Catalog no. 00-4956-58) was performed after deparaffinization on 8 mm-thick tissue sections. Endogenous peroxidases were quenched with 3% H2O2 in methanol for 30 minutes. The sections were blocked in 5% normal goat serum at room temperature for 1 hour, incubated with primary antibody overnight at 4°C and secondary antibody at room temperature for 1 hour, followed by color development in DAB using Metal Enhanced DAB Substrate Kit (ThermoFisher Scientific, Catalog no. 34065).

### Gel Electrophoresis and Western Blotting

For immunoblotting in Fig. 3 and 4, liver or tumor protein lysates were prepared in radioimmune precipitation assay buffer with proteinase inhibitors, and protein concentrations of lysates were determined using a Bradford protein assay reagent (Bio-Rad). Each sample was resolved by electrophoresis using 10% SDS-PAGE and transferred to PVDF membranes. After blocking with 5% dry milk, analysis of proteins was performed using antibodies against CYP2E1 (Invitrogen, Catalog no. MA5-32605), glyceraldehyde-3-phosphate dehydrogenase (GAPDH) (Invitrogen, Catalog no. AM4300) and Zhx2 (Proteintech, Catalog no. 20136-1-AP) and SuperSignal™ West Pico PLUS Chemiluminescent Substrate kit (ThermoFisher Scientific, Catalog no. 34577). The blots were visualized using Amersham Imager 600 (GE Healthcare) and the protein levels were quantified using ImageQuant™ TL Image Analysis Software (GE Healthcare).

For Western blotting in Fig. 5, ∼50 mg of liver tissue was resuspended in 3 volumes of M-PER Mammalian Protein Extraction Reagent (ThermoFisher Scientific, Cat no: 78501) plus 10% protease inhibitor cocktail (Sigma P2714-1BTL) and Halt phosphatase inhibitor cocktail (Fisher PI78420), and incubated on ice for 30 min. The samples were lysed using a Qiagen Tissue Lyser LT (Qiagen) and were resolved by SDS polyacrylamide gel electrophoresis and electrophoretically transferred to Immobilon-FL membranes. Membranes were blocked at room temperature for 1 h in the LI-COR Intercept Blocking Buffer (LI-COR Biosciences, Cat no: 927-60001). Subsequently, the membranes were incubated overnight at 4°C with the following antibodies: Serine 473 phospho-AKT (Cell Signaling, Cat no: 4060S), AKT1/2 (Santa Cruz Biotechnology, Cat no: sc-1619), AKT1 (Santa Cruz Biotechnology, Cat no: sc-5298), AKT2 (Santa Cruz Biotechnology, Cat no: sc-5270), NF-κB p65 (Abcam, Cat no: ab31481),or heat shock protein 90 (HSP90) (Santa Cruz Biotechnology, Cat no: sc-13119). After three washes, the membrane was incubated with an infrared anti-rabbit (IRDye 800, green) or anti-mouse (IRDye 680, red) secondary antibody labeled with IRDye infrared dye (LI-COR Biosciences) for 2 h at 4°C. Immunoreactivity was visualized and quantified by infrared scanning in the Odyssey system (LI-COR Biosciences).

**Supplementary Table 1.**
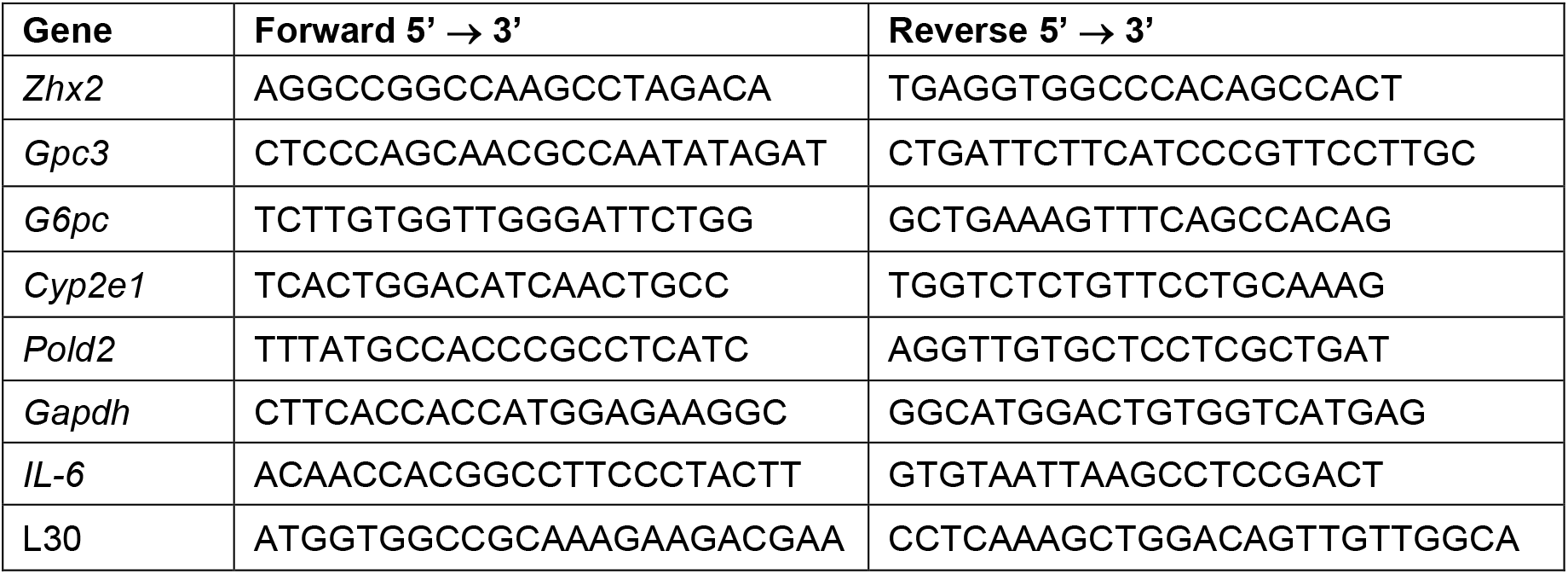
Oligonucleotides used for RT-qPCR.

**Supplementary Table 2.**
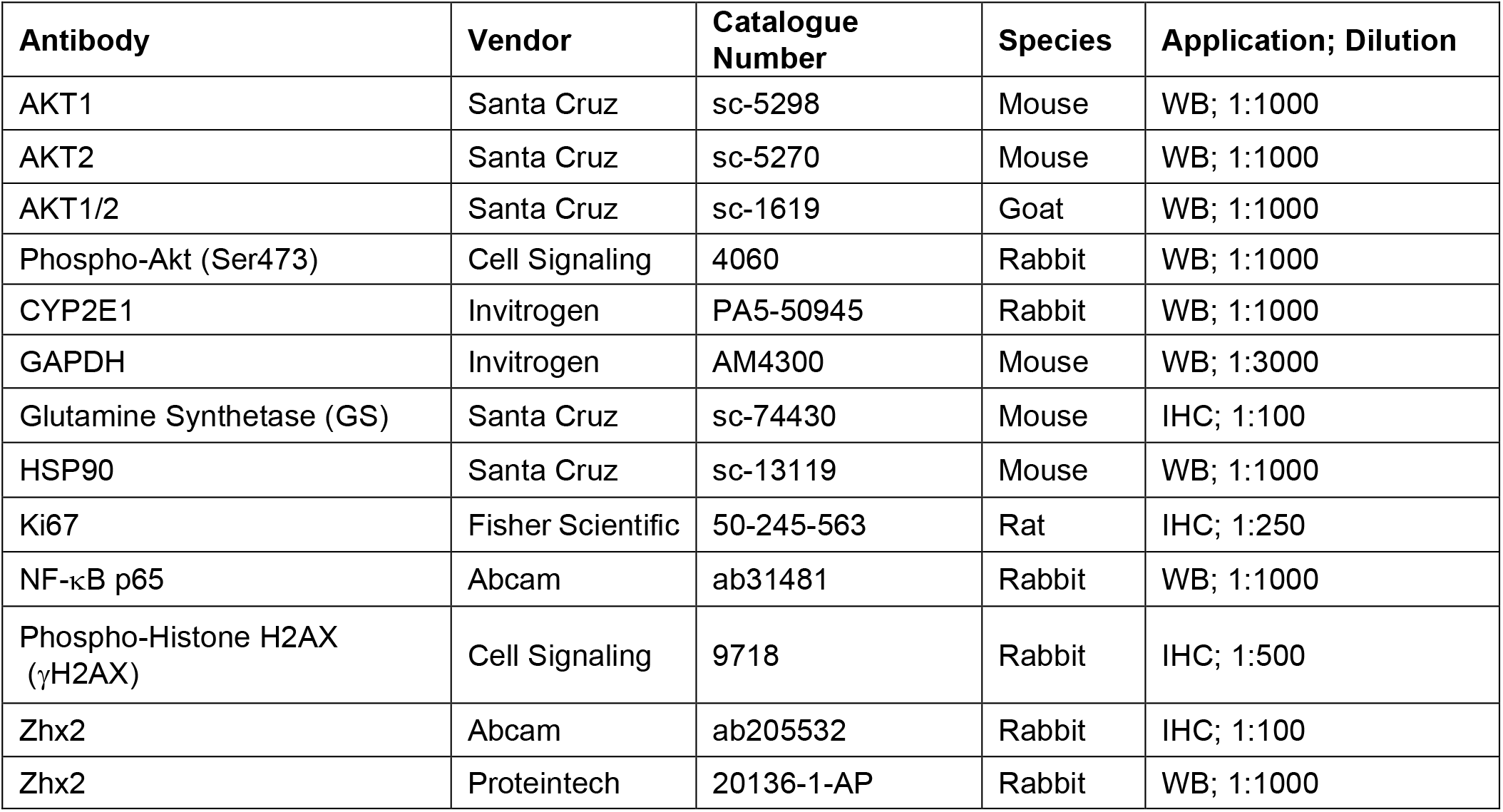
Antibodies used in this study.

**Supplementary Figure 1.**
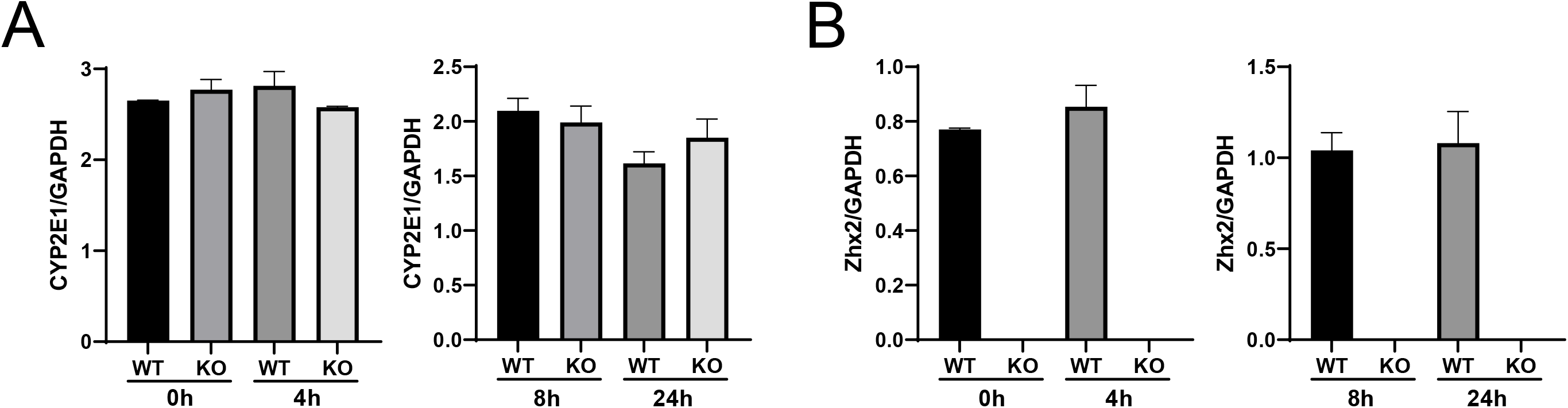
Quantification of CYP2E1 and Zhx2 protein expression in *Zhx2*^*wt*^ and *Zhx2*^*KO*^ mice 24 hours after DEN treatment. Hepatic CYP2E1 and Zhx2 protein levels from p14 *Zhx2*^*wt*^ and *Zhx2*^*KO*^ mice prior to (0h) and 4, 8, or 24 hours after DEN treatment were analyzed by Western blot as shown in Figure 4 C, D. The CYP2E1 (**A**) and Zhx2 (**B**) levels were quantified using ImageQuant™ TL Image Analysis Software (GE Healthcare) and normalized to GAPDH.

